# The maternal microbiome promotes placental development in mice

**DOI:** 10.1101/2023.02.15.528712

**Authors:** Geoffrey N. Pronovost, Sahil S. Telang, Angela S. Chen, Elena J.L. Coley, Helen E. Vuong, Drake W. Williams, Kristie B. Yu, Tomiko K. Rendon, Jorge Paramo, Reuben H. Kim, Elaine Y. Hsiao

## Abstract

The maternal microbiome is an important regulator of gestational health, but how it impacts the placenta as the interface between mother and fetus remains unexplored. Here we show that the maternal gut microbiota supports placental development in mice. Depletion of the maternal gut microbiota restricts placental growth and impairs feto-placental vascularization. The maternal gut microbiota modulates metabolites in the maternal and fetal circulation. Short-chain fatty acids (SCFAs) stimulate angiogenesis-related tube formation by endothelial cells and prevent abnormalities in placental vascularization in microbiota-deficient mice. Furthermore, in a model of maternal malnutrition, gestational supplementation with SCFAs prevents placental growth restriction and vascular insufficiency. These findings highlight the importance of host-microbial symbioses during pregnancy and reveal that the maternal gut microbiome promotes placental growth and vascularization in mice.

## Main Text

Recent studies highlight notable influences of the maternal microbiome on offspring development that begin during the prenatal period (*1, 2*), but exactly how the maternal microbiome informs maternal-fetal health during pregnancy remains unclear. At the intersection of mother and fetus is the highly vascularized placenta, which is responsible for enabling maternal-fetal exchange of nutrients and gases that sustain fetal development (*3, 4*). We examined effects of the maternal gut microbiome on the development of the placenta, as a critical organ that shapes long-term health trajectories.

To determine effects of the maternal gut microbiome on placental development, we first reared pregnant mice as germ-free (GF) or depleted the maternal gut microbiome by treating with broadspectrum antibiotics (ABX). Absence or depletion of the maternal gut microbiome resulted in reduced placental weight at embryonic day 0.5 (E0.5), relative to conventionally colonized (specific pathogen-free, SPF) and conventionalized GF (GF CONV) controls (Fig. 1, A and B). Maternal treatment with the subset of ABX that are non-absorbable phenocopied the reduced placental weight seen with the full ABX cocktail (fig. S1, A to D), with expected reductions in microbial diversity (fig. S1, E to G). This suggests that ABX-induced reductions in placental weight are due to depletion of the maternal gut microbiome rather than off-target effects of ABX. To determine if changes in placental weight are due to localized disruptions to particular subregions of the placenta, placentas from gnotobiotic dams were imaged by microcomputed tomography (*μ*CT). Consistent with reductions in placental weight, maternal microbiome deficiency led to reductions in total placental volume and volume of the placental labyrinth, in particular, as the primary site for maternal-fetal exchange (Fig. 1, C to E). In addition to maternal ABX-induced placental pathophysiology, we observed corresponding decreases in fetal weight and volume (fig. S2) which were not seen in the GF condition. Overall, these data reveal a key role for the maternal gut microbiome in promoting placental development, particularly within the labyrinth subregion.

**Fig. 1:**
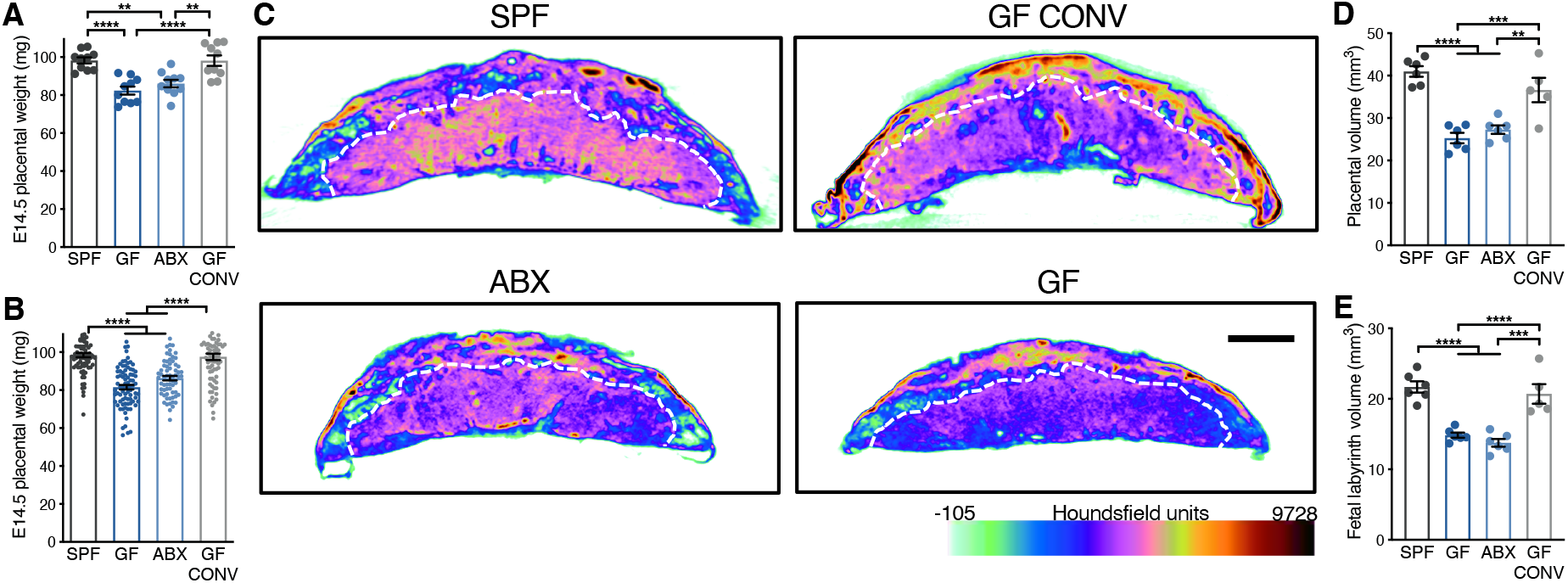
The maternal microbiome promotes placental development. (**A)** E14.5 placental weights by litter average (SPF (*n* = 10), GF (*n* = 10), ABX (*n* = 10) and GF CONV (*n* = 10). (**B)** E14.5 placental weights for each individual from litters shown in Fig. 1A (SPF (*n* = 80), GF (*n* = 81), ABX (*n* = 65) and GF CONV (*n* = 77). (**C)** Representative cross-sections of E14.5 whole placental *μ*CT reconstructions from SPF, GF, ABX and GF CONV litters. White hashed line distinguishes fetal placental labyrinth compartment; scale bar = 1 mm; Houndsfield scale range from −105 (light green) to 9728 (maroon). (**D)** Quantification of E14.5 whole placental volumes from *μ*CT reconstructions by litter average (SPF (*n* = 6), GF (*n* = 6), ABX (*n* = 6) and GF CONV (*n* = 5)). (**E)** Quantification of fetal placental labyrinth volumes from *μ*CT reconstructions shown in Fig. 1D. Data represent mean ± SEM; statistics were performed with one-way ANOVA with Tukey *post hoc* test. \**P* < 0.05; \*\**P* < 0.01; \*\*\**P* < 0.001; \*\*\*\**P* < 0.0001.

The placental labyrinth is comprised of maternal and fetal blood spaces separated by trophoblast cells, basement membrane, and fetal endothelial cells that together direct gas and nutrient exchange. To gain insight into potential cellular bases underlying the placental deficits induced by maternal microbiome deficiency, we performed RNA sequencing on microdissected labyrinth tissues. Absence or depletion of the maternal gut microbiome led to shared differential regulation of 57 genes (Fig. 2A), with many of the downregulated genes relating to pathways important for placental vascular development (*5, 6*). This led us to hypothesize that maternal microbiome-induced changes in placental labyrinth gene expression could reflect altered development of fetoplacental vasculature. To test this, we stained placentas with the endothelial cell marker laminin, and saw significantly reduced staining intensity in placental labyrinth tissue from microbiome deficient dams relative to controls (Fig. 2, B to C). We further generated feto-placental arterial casts and analyzed their native structures using *μ*CT imaging (*13*). Feto-placental vasculature from microbiota deficient dams exhibited reduced vascular volume and surface area, with visible decreases in vascular branching, as compared to controls (Fig. 2, D to F). This aligns with prior studies that have reported a role for the microbiome in regulating angiogenesis and barrier integrity in other organs, including the intestine and brain (*7, 8*). Collectively, these data reveal that the maternal microbiome is required for proper development of feto-placental vasculature.

**Fig. 2:**
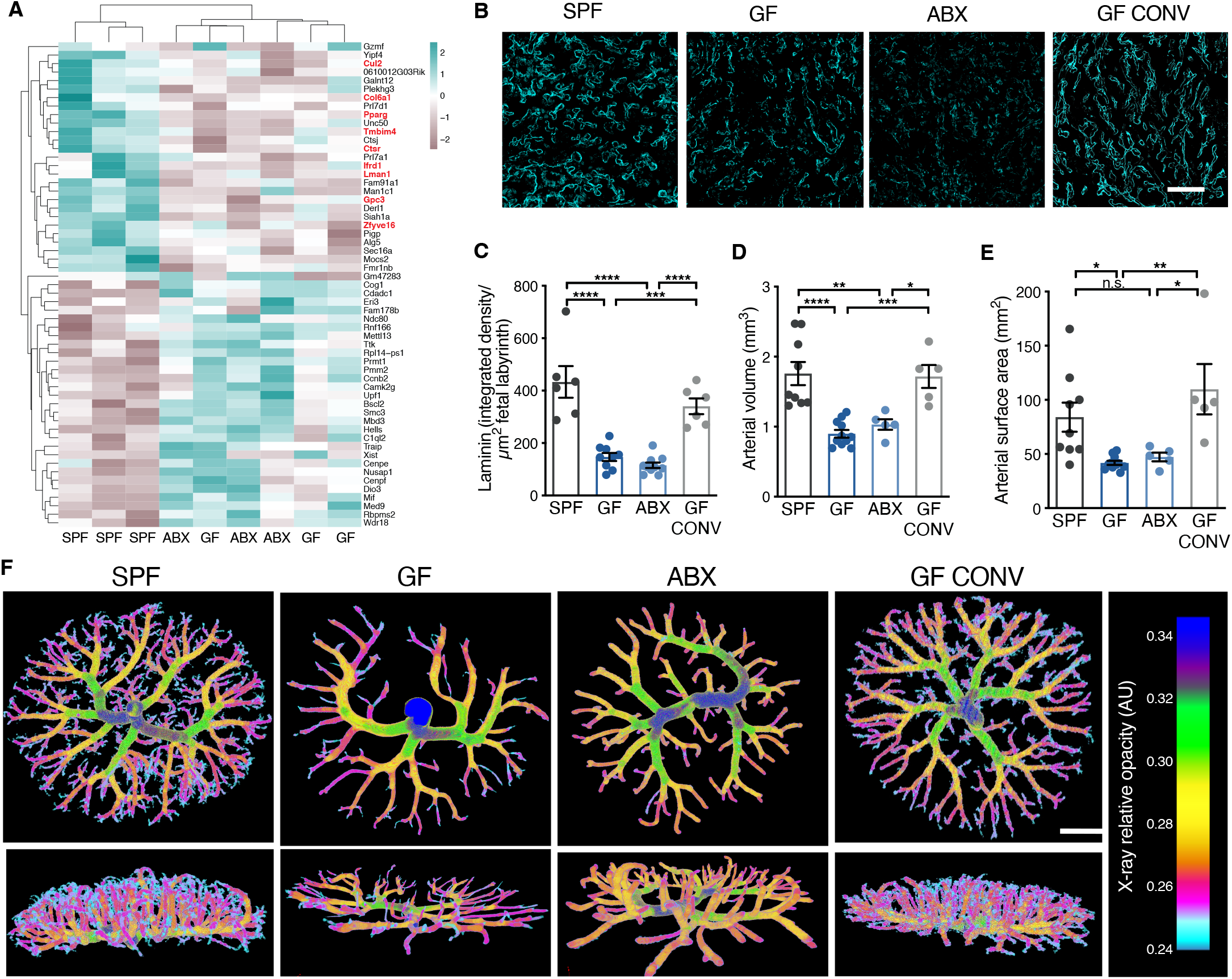
The maternal microbiome promotes placental vascular development. **(A)** Significantly differentially expressed genes (two-tailed Wald, *P*<0.05) from E14.5 fetal labyrinth (SPF (*n* = 3), GF (*n* = 3) and ABX (*n* = 3)); vascular-related genes labeled in red text. (**B)** Representative images of laminin-stained E14.5 placental labyrinth from SPF, GF, ABX and GFCONV litters (scale bar = 50 μm). (**C)** Quantification of raw integrated density of laminin staining intensity, normalized to fetal labyrinth total area (SPF (*n* = 6), GF (*n* = 9), ABX (*n* = 10) and GF CONV (*n* = 5)). (**D)** Quantification of E14.5 feto-placental arterial vascular volume (SPF (*n* = 9), GF (*n* = 11), ABX (*n* = 5) and GF CONV (*n* = 5)). (**E)** Quantification of E14.5 feto-placental arterial vascular surface area (SPF (*n* = 9), GF (*n* = 11), ABX (*n* = 5) and GF CONV (*n* = 5)). (**F)**. Representative feto-placental arterial vascular reconstructions by *μ*CT imaging of vascular casts from E14.5 SPF, GF, ABX and GF CONV litters; scale bar = 1 mm. Data represent mean ± SEM; statistics were performed with one-way ANOVA with Tukey *post hoc* test. \**P* < 0.05; \*\**P* < 0.01; \*\*\**P* < 0.001; \*\*\*\**P* < 0.0001.

Given that the maternal microbiome regulates many circulating metabolites (*2*), we postulated that microbiome-dependent insufficiencies in feto-placental vasculature could arise from alterations in key metabolites in the fetal circulation. Therefore, we performed untargeted metabolomic profiling of 753 metabolites in the fetal serum metabolome (Table 1). By principal components analysis, fetal serum metabolomic profiles from GF and ABX dams generally clustered away from SPF controls (fig. S3A). 27 metabolites were commonly and significantly downregulated and 14 were commonly and significantly upregulated in fetal serum from GF and ABX dams, compared to SPF controls (fig. S3, B to D). Random forest analysis identified 30 fetal serum metabolites that predicted maternal microbiota status with 89% accuracy (fig. S3E). No statistically significant changes were seen for fetal serum metabolites related to core metabolism, including glucose and many amino acids (Table 1), suggesting that the placental and fetal deficits caused by maternal microbiome deficiency are not due to overt fetal nutrient restriction. These data indicate that the maternal microbiome modulates the bioavailability of metabolites in the fetal circulation, which is consistent with previous data demonstrating that the maternal microbiome regulates metabolites in the maternal serum and fetal brain during pregnancy (*2*). To determine whether select maternal microbiome-dependent metabolites in the fetal serum promote feto-placental vascular development and placental growth, we supplemented microbiota-deficient dams with a pool of 19 metabolites that were significantly and commonly reduced in both maternal serum and fetal serum from microbiota-deficient dams (fig. S3, F to J). Daily systemic injection of the microbially modulated metabolites at physiologically-relevant concentrations failed to prevent maternal ABX-induced reductions in placental and fetal weight (fig. S4, A to C) and in microvascular laminin staining of placental tissues (fig. S4, D and E). Thus, the placental insufficiencies induced by maternal microbiome deficiency are likely not regulated by this select group of maternal microbiome-dependent metabolites.

In addition to the metabolites identified and tested, we considered the role that microbial generation of short-chain fatty acids (SCFAs) might play in regulating placental and fetal development (*9–11*). SCFAs are produced by bacterial carbohydrate fermentation and are significantly decreased in the maternal and fetal serum from microbiota-deficient dams (*12*). Based on previous research demonstrating that maternal supplementation with SCFAs leads to direct transfer of SCFAs from maternal circulation to fetal circulation (*12*), we treated ABX dams with SCFA-supplemented water (*11*) or vehicle (sodium-matched) control water from E0.5 to E14.5. Maternal SCFA treatment increased placental weight from ABX dams to levels comparable to SPF controls, with corresponding increases in total placental and labyrinth volumes as measured by *μ*CT imaging (Fig. 3, A to D). Maternal SCFA treatment elicited a modest increase in fetal weights from microbiota-depleted dams, which reached statistical significance when considering all individuals across litters, but not litter averages (fig. S5). Consistent with the increases in placental growth, maternal SCFA treatment increased endothelial laminin staining in placental labyrinth from ABX dams toward levels seen in SPF controls (Fig. 3, E and F). These observations were further supported using *μ*CT-reconstructions of feto-placental arterial casts, where maternal SCFA treatment increased fetal vascular volume and surface area in placentas from ABX dams (Fig. 3, G to I). These data reveal that microbiota-derived SCFAs promote placental growth and fetoplacental vascular development.

**Fig. 3:**
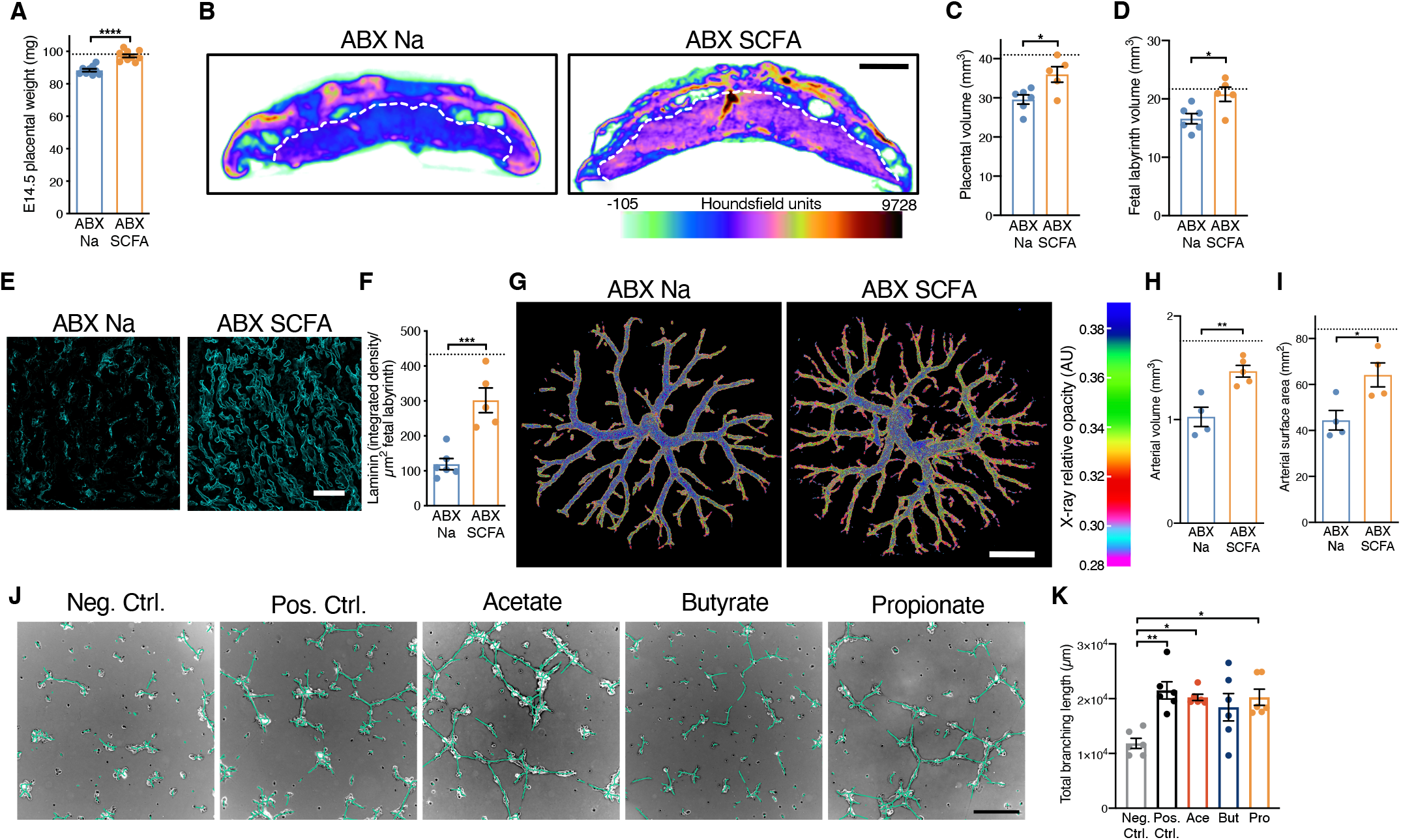
Maternal SCFA supplementation prevents impairments in placental growth and microvasculature induced by maternal microbiome depletion. **(A)** E14.5 placental weights by litter average (ABX Na (*n* = 9) and ABX SCFA (*n* = 10); hashed line represents SPF litter average value shown in Fig. 1A). (**B)** Representative cross-sections of E14.5 whole placental *μ*CT reconstructions from ABX Na and ABX SCFA litters. (**C)** Quantification of E14.5 whole placental volumes from *μ*CT reconstructions by litter average (ABX Na (*n* = 6) and ABX SCFA (*n* = 5); hashed line represents SPF litter average value shown in Fig. 1D). (**D)** Quantification of fetal placental labyrinth volumes from *μ*CT reconstructions shown in Fig. 3D; hashed line represents SPF litter average value shown in Fig. 1E. (**E)** Representative images of laminin-stained E14.5 placental labyrinth from ABX Na and ABX SCFA litters; scale bar = 50 μm. (**F)** Quantification of raw integrated density of laminin staining intensity, normalized to fetal labyrinth total area (ABX Na (*n* = 6), ABX SCFA (*n* = 5); hashed line represents SPF litter average value shown in Fig. 2C). (**G)** Representative feto-placental arterial vascular reconstructions by *μ*CT imaging of vascular casts from E14.5 ABX Na and ABX SCFA litters; scale bar = 1 mm. (**H)** Quantification of E14.5 feto-placental arterial vascular volume (ABX Na (*n* = 4) and ABX SCFA (*n* = 4); hashed line represents SPF litter average value shown in Fig. 2D). (**I)** Quantification of E14.5 feto-placental arterial vascular surface area from litters shown in Fig. 3H; hashed line represents SPF litter average value shown in Fig. 2E. (**J)** Representative images of HUVEC tube formation assays (scale bar = 250 μm), depicting negative control (no supplementation), positive control (4% FBS supplementation), acetate supplementation (Ace, 40 μm), butyrate supplementation (But, 5 μm) and propionate supplementation (Pro, 5 μm). (**K)** Quantification of HUVEC tube formation assays from independent experiments (*n* = 6). Data represent mean ± SEM; statistics were performed with student’s t test or one-way ANOVA with Tukey *post hoc* test. \**P* < 0.05; \*\**P* < 0.01; \*\*\**P* < 0.001; \*\*\*\**P* < 0.0001.

SCFAs have been reported to protect against aortic endothelial dysfunction and promote fibrovascular angiogenesis through direct receptor-mediated signaling (*13, 14*). To gain insight into whether SCFAs may signal directly to fetal endothelial cells to promote vascularization, we treated human umbilical vein endothelial cells (HUVECs) with SCFAs at physiological concentrations (*15, 16*) and quantified tube formation as a measure of angiogenesis. The SCFAs acetate and propionate significantly increased HUVEC branching length compared to vehicle controls, whereas butyrate elicited variable increases in branching length that did not meet statistical significance (Fig. 3, J and K, fig. S6). Increased tube formation was not seen when cells were treated with various other microbiome-dependent metabolites (fig. S7), which were included in the pool that failed to prevent placental disruptions when administered to ABX dams (Fig. 3), suggesting specificity to SCFAs. Overall, these data indicate that SCFAs directly stimulate vascularization of cultured umbilical vein endothelial cells, consistent with our findings that SCFA supplementation promotes placental vascular development *in vivo*.

Maternal malnutrition, including maternal protein restriction (PR), is associated with placental insufficiencies, including reduced size and impaired vasculature (*17, 18*). Based on the ability of the maternal microbiome and SCFAs to promote placental development in dams with deficient microbiomes (Figs. 1 to 3), we next asked whether maternal SCFA treatment could prevent placental abnormalities caused by maternal PR during pregnancy. To isolate effects of PR to restrictions in protein, rather than compensatory increases in complex carbohydrates, the PR diet and a control diet (CD) with levels of protein matching those in standard mouse chow were formulated with cellulose as the primary carbohydrate source. Consistent with previous findings (*18*), maternal PR reduced placental weight and volume, when compared to dams fed CD (Fig. 4, A to D). Maternal SCFA treatment was sufficient to restore placental weight, total volume and labyrinth-specific volume in litters from PR dams to levels seen in CD and previous SPF controls (Fig. 4, A to D). Consistent with its use as a model of intrauterine growth restriction, maternal PR induced modest decreases in fetal weight, which were statistically significant when considering individual conceptuses but not litter averages (fig. S8). These reductions in fetal weight were prevented by maternal SCFA treatment of PR dams (fig. S8). Moreover, maternal SCFA treatment increased feto-placental vascularization, as measured by increases in feto-placental arterial vascular volume and surface area (Fig. 4, E to G). Taken together, these data suggest that SCFAs promote placental growth and vascular development, even under conditions of maternal malnutrition.

**Fig. 4:**
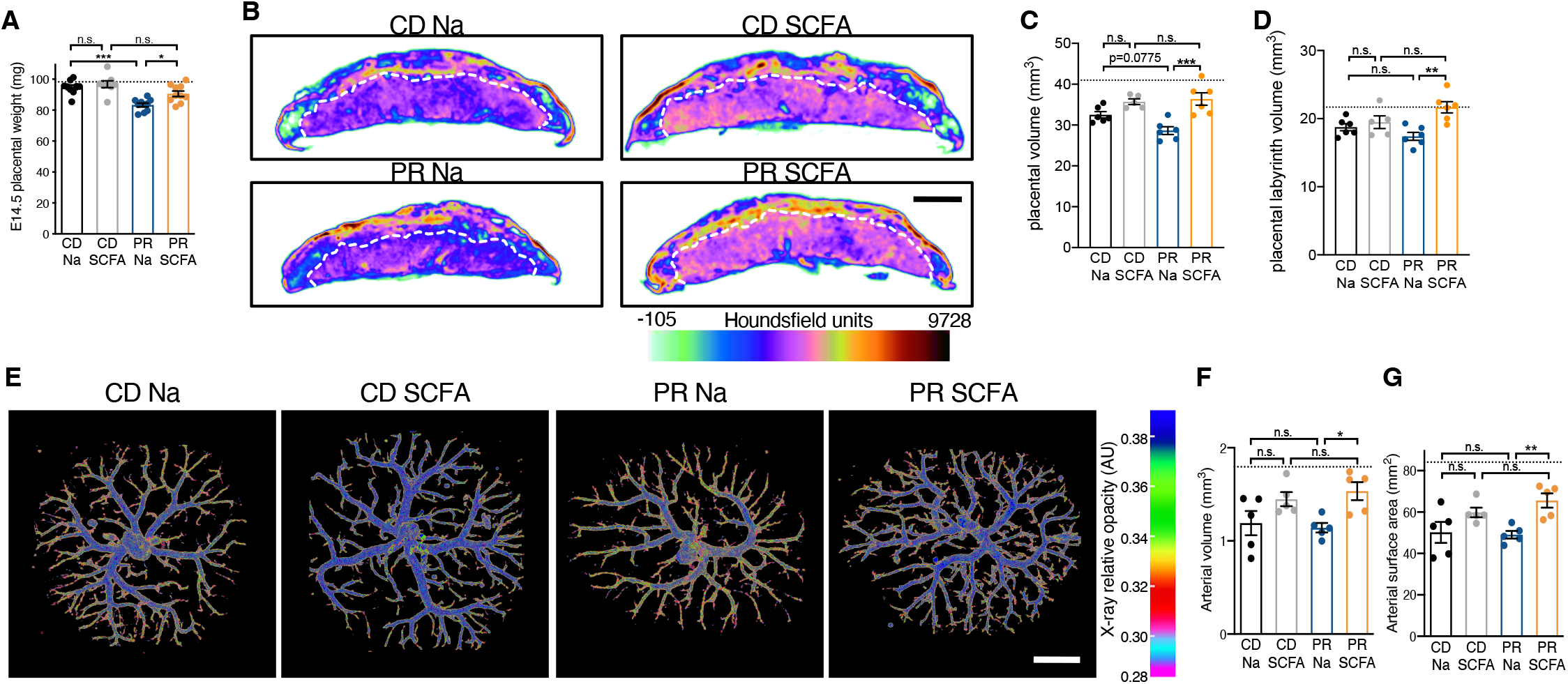
Maternal SCFA supplementation promotes placental growth and vascularization in protein-restricted dams. **(A)** E14.5 placental weights by litter average (CD Na (*n* = 9), CD SCFA (*n* = 10), PR Na (*n* = 8) and PR SCFA (*n* = 10); hashed line represents SPF litter average value shown in Fig. 1A). (**B)** Representative cross-sections of E14.5 whole placental *μ*CT reconstructions from CD Na, CD SCFA, PR Na and PR SCFA litters. (**C)** Quantification of E14.5 whole placental volumes from *μ*CT reconstructions by litter average (CD Na (*n* = 6), CD SCFA (*n* = 5), PR Na (*n* = 6) and PR SCFA (*n* = 5); hashed line represents SPF litter average value shown in Fig. 1D). (**D)** Quantification of fetal placental labyrinth volumes from *μ*CT reconstructions shown in Fig. 4C; hashed line represents SPF litter average value shown in Fig. 1E. (**E)** Representative feto-placental arterial vascular reconstructions by *μ*CT imaging of vascular casts from CD Na, CD SCFA, PR Na and PR SCFA litters; scale bar = 1 mm. (**F)** Quantification of E14.5 feto-placental arterial vascular volume (CD Na (*n* = 5), CD SCFA (*n* = 5), PR Na (*n* = 5) and PR SCFA (*n* = 5); hashed line represents SPF litter average value shown in Fig. 2D). (**G)** Quantification of E14.5 feto-placental arterial surface area from litters shown in Fig. 4F; hashed line represents SPF litter average value shown in Fig. 2D). Data represent mean ± SEM; statistics were performed with two-way ANOVA with Tukey *post hoc* test. *\*P* < 0.05; *\*\*P* < 0.01; *\*\*\*P* < 0.001; *\*\*\*\*P* < 0.0001.

Results from this study reveal that the maternal microbiome promotes placental development in mice and that specific byproducts of bacterial metabolism, SCFAs, instruct placental growth and vascularization. Maternal treatment with SCFAs during pregnancy prevents impairments in feto-placental development that are caused by maternal microbiome deficiency and maternal protein undernutrition. These results open the possibility that microbiome-directed interventions may be leveraged to promote feto-placental health during pregnancy.

## Acknowledgments

We thank S. Devaskar, L. Iruela-Arispe, and all members of the Hsiao lab for their helpful feedback on the study.

## Author contributions

Conceptualization: GNP, EYH

Methodology: GNP, SST, ASC, EJC, KBY, HEV, DWW

Funding acquisition: GNP, EYH

Resources: RHK, EYH

Writing – original draft: GNP, EYH

Writing – review & editing: GNP, SST, ASC, EJC, KBY, HEV, DWW, RHK, EYH

## Data and materials availability

Data generated or analyzed during this study are included in this published article and its supplementary information files. The 16S rRNA gene sequencing data that support the findings of this study are available in Table S4 and have been deposited to the Qiita database with study ID 14433. Transcriptomic data that support the findings of this study are available in the Table S5 and have been deposited to the Gene Expression Omnibus (GEO) repository (GSE198947).

## Supplementary Materials

Materials and Methods

Supplementary Text

Fig. S1 – Fig. S8

References (19-32

## Materials and Methods

### Mice

C57Bl/6J mice were purchased from Jackson Laboratories and either reared as SPF, or rederived as GF and bred in flexible film isolators at the UCLA Center for Health Sciences barrier facility. Animals were maintained on a 12-h light-dark cycle in a temperature-controlled environment, with autoclaved bedding. Sterile water and autoclaved standard chow (Lab Diet 5010) were provided *ad libitum*. For all experiments, pregnant dams were sacrificed by cervical dislocation to preclude the effects of CO_2_ exposure on maternal and fetal physiology. All experiments were performed in accordance with the NIH Guide for the Care and Use of Laboratory Animals using protocols (2015-077-01) approved by the Institutional Animal Care and Use Committee at UCLA.

#### Sample size determination

8-week-old mice were randomly assigned to experimental groups, which included age- and sex-matched cohorts of males and females for timed mating. To account for maternal microbiome status as the primary experimental variable in all animal experiments, biological sample sizes reflect the number of independent dams, and only virgin females were used for breeding. Characterization of placental and fetal phenotypes included at least 2 randomly selected conceptuses per litter for each individual dam, where litter averages represent biological “n”. For placental and fetal weight statistics, litter averages are presented in the main figures of the manuscript, and accompanying individual conceptus data are included either in main figures or in supplementary figures.

#### Antibiotic treatment and conventionalization

For broad-spectrum antibiotic treatment (ABX), 8 week-old SPF mice were gavaged twice daily (08:00 and 17:00) for 1 week with a cocktail of neomycin (100 mg/kg), metronidazole (100 mg/kg), and vancomycin (50 mg/kg), based on methods previously described to mimic GF status (*1*). Ampicillin (1 mg/ml) was provided *ad libitum* in drinking water. Breeders were then paired and time-mated. Embryonic day (E) 0.5 was determined by observation of the copulation plug using a sterile vaginal probe. Dams were then separated, individually housed and maintained on sterile drinking water supplemented with ampicillin (1 mg/ml), neomycin (1 mg/ml) and vancomycin (0.5 mg/ml) until E14.5 to preclude any stress of oral gavage in pregnant dams.

For selective antibiotic treatment, 8-week-old SPF mice were randomly selected for treatment with either antibiotics that are absorbed into host circulation (Abs) or with antibiotics that are not absorbed into host circulation (Non-Abs). Abs mice were gavaged twice daily (08:00 and 17:00) for 1 week with metronidazole (100 mg/kg), and ampicillin (1 mg/ml) was provided *ad libitum* in drinking water. Non-Abs mice were gavaged twice daily (08:00 and 17:00) for 1 week with neomycin (100 mg/kg) and vancomycin (50 mg/kg), and then maintained on sterile drinking water with neomycin (1 mg/ml) and vancomycin (0.5 mg/ml). Breeders were paired and time-mated, and dams were separated on E0.5 as described above.

#### Conventionalization of GF mice

SPF donor fecal pellets were freshly homogenized at 100 mg/ml in pre-reduced sterile PBS, and 200 μl was gavaged into each 8-week-old female GF recipients once per week for 2 weeks. Bedding from the SPF donor home cage was also added to the GF CONV recipient cage to maximize conventionalization.

#### Maternal protein-restriction

8-week-old mice previously fed standard chow (Lab Diet 5010) were randomly selected to receive either control diet with 20.3% protein by weight, 61.3% carbohydrate by weight and 5.5% fat by weight (TD.91352, Envigo), or protein-restricted diet with 6.1% protein by weight, 75.6% carbohydrate by weight and 5.5% fat by weight (TD.90016, Envigo). Differences in carbohydrates resulted from amounts of sucrose and cellulose, and both diets used were isocaloric by weight (3.8 Kcal/g) and matched for calcium (0.7%) and phosphorous (0.54%). Both male and female mice acclimated to the respective diets for a 2-week period and were then paired for breeding within the respective dietary treatment groups. Upon observation of the copulation plug at E0.5, females were individually housed and maintained on the respective diet for the duration of gestation.

### 16S rRNA gene sequencing

Bacterial genomic DNA was isolated from mouse fecal samples following the standard protocol from the DNeasy PowerSoil Kit (Qiagen). Sequencing libraries were generated according to methods adapted from Caporaso et al. 2011 (*2*), amplifying the V4 regions of the 16S rRNA gene by PCR using individually barcoded universal primers and 30 ng of the extracted genomic DNA. Each PCR reaction was performed in triplicate and pooled following amplification. The final PCR product was purified using the Qiaquick PCR purification kit (Qiagen). 250 ng of purified, PCR product from each individually barcoded sample were pooled and sequenced by Laragen, Inc. using the Illumina MiSeq platform and 2 x 250bp reagent kit for paired-end sequencing. All analyses were performed using QIIME2 (*3*), including Deblur for quality control (*4*), taxonomy assignment, alpha-rarefaction and beta-diversity analyses. 29,152 reads were analyzed per sample. Operational taxonomic units (OTUs) were assigned based on 99% sequence similarity compared to the SILVA 132 database. Alpha-rarefaction curves and beta-diversity principal coordinates analysis plots were generated using Prism software (GraphPad) and QIIME2 View, respectively, and taxa bar plots were generated using Microsoft Excel.

### Placental RNA sequencing

Dams were sacrificed on E14.5 by cervical dislocation, and uterine horns were dissected immediately and placed in ice-cold PBS. Two randomly chosen conceptuses per dam were isolated from the uterine horn. Placentas were cut from the base of the umbilical cord, micro-dissected to remove the underlying fetal placental tissue, pooled and collected into RNA*later* (ThermoFisher Scientific), and frozen at −80°C. Frozen tissues were thawed, placed in Trizol (Invitrogen), maintained on ice and homogenized using 5 mm stainless steel beads (Qiagen) on a minibeadbeater (BioSpec) for 7 seconds. RNA was extracted using the RNAeasy Mini kit with on-column genomic DNA-digest (Qiagen), and RNA quality of RIN > 8.8 was confirmed using the 4200 Tapestation system (Agilent). RNA was prepared using the TruSeq RNA Library Prep kit and Lexogen QuantSeq 3’ forward-sequencing was performed using the Illumina HiSeq 4000 platform by the UCLA Neuroscience Genomics Core. Quality filtering and mapping were performed using the BlueBee analysis platform (Lexogen), and reads were aligned to the UCSC Genome Browser assembly ID:mm10. At least 1.07 million aligned reads were obtained for each sample. Differential gene expression analysis was conducted using DESeq2 1.24.0 (*5*), and genes of interest were selected based on p < 0.05.

### Immunofluorescence staining

E14.5 placentas were fixed in 4% paraformaldehyde for 24 hours at 4°C, cryoprotected in 30% sucrose in PBS for 24 hours at 4°C, and sectioned at 12 μm using a Leica CM1950 cryostat. Sections were blocked with 10% goat serum for 1 hour at room temperature. Primary antibodies were diluted in 10% goat serum and incubated for 18 hours at 4°C with laminin rabbit anti-mouse antibody (1:250, Sigma-Aldrich, L9393). Sections were then incubated for 1 hour at room temperature in their corresponding goat anti-rabbit antibodies conjugated to Alexa Fluor 568 (1:1000, Thermofisher Scientific) and DAPI. Images were acquired using the Zeiss Axio Examiner LSM 780 confocal microscope with 405nm (0.2% laser line attenuator transmission, 575 master gain, 0 digital offset, 1.0 digital gain) and 561nm (2.0% laser line attenuator transmission, 605 master gain, −2.0 digital offset, 1.0 digital gain) lasers. 1-2 placental sections were scanned for each sample using Zen Black 2012 software, 20X objective at 1.5X zoom with 5 x 1μm interval z-slices and 3 individual tracks for each fluorescent dye. Image acquisition settings included: scan mode set at frame, frame size set at 1024 X 1024, scan speed set at 7, averaging at 2 by line and mean, and bit depth set at 8 bit. Pinhole was set to 1AU. Images were stitched with a 10%-overlap, and the complete range of z-series were compressed using Zen Blue 2021 software.

#### Image analysis

All acquired images were analyzed using the same procedures by a researcher blinded to the experimental group of each sample. Images were imported into Fiji and calibrated using a set scale. The perimeter of the labyrinth area was manually traced, and labyrinth area was calculated by oversaturating all tissue using brightness and contrast settings from 0 minimum to 8 maximum, where areas of no signal were defined as void spaces. To quantify integrated density of laminin, brightness and contrast settings were adjusted to eliminate background auto-fluorescence. Raw integrated density values for each sample were then normalized to the total labyrinth area for each sample.

### Microcomputed tomography

Whole E14.5 placentas and fetuses were serially dehydrated, from 30%- to 50%- to 70% ethanol and incubated in 4% (w/v) phosphotungstic acid (EPTA) diluted in 70% ethanol for 4 days at 4°C. Tissues were scanned at 60 kVp/150 μA with a 1 mm Al filter at 8 μm resolution using a benchtop *μ*CT scanner (SkyScan 1275, Bruker). Reconstructions of 2-dimensional images were generated using dynamic range adjustment and gaussian smoothing, a ring artifact reduction of 10, and a defect pixel mask of 8%. Volumes of interest (VOI) for placental labyrinth volume measurements were selected manually. Whole placental volumes were reconstructed and measured using CTAn and CTVol software (Bruker Corporation), using a threshold range of 88 (minimum) to 255 (maximum). Whole placenta representative images were created using Dataviewer software (Bruker Corporation), with heat maps reflecting Houndsfield units as a metric of radio density.

#### Placental arterial vascular casts

Feto-placental arterial vascular casts were generated adapting previously described methods (*6*). Briefly, dams were sacrificed on E14.5 by cervical dislocation, and uterine horns were dissected immediately and placed in ice-cold PBS. Conceptuses were individually removed, placed on a heating pad set at 37°C and continuously flushed with warmed (37°C) PBS to restore fetal heartbeat. The fetal umbilical artery was identified following observation of pulsing blood flow, 4% PFA in PBS was dropped onto the artery for partial vasodilation, and a micro-incision was made on the artery. A hand-pulled glass pipette affixed to tubing was inserted into the arterial lumen, and silicone adhesive KwikSil (World Precision Instruments) was used to cover the incision site and adhere the pipette in place. Once the silicone adhesive was set, a small incision was made on the umbilical vein to serve as an outflow for perfusions. To flush the feto-placental vasculature and ensure equal vasodilation across all samples, a warmed (37°C) 1:1 solution of heparinized (100 Iu/ml) PBS and 10% lidocaine was perfused through the placenta until the perfusate was free of blood. Without introducing air bubbles, a tubing line with ice-cold MICROFIL silicone rubber injection compound (Flow Tech Inc.) was perfused through the umbilical artery until resistance was met to maintain pressure. Both umbilical vessels were immediately sutured and filled placentas were placed in room temperature PBS for at least 30 minutes for casting compound to polymerize. Samples were then serially dehydrated in ethanol, from 30%-, 50%- into 70%-ethanol at 4°C to reduce background electron density from placental tissue. Perfused placental casts were imaged by microcomputed tomography, as described above. Feto-placental arterial volume and surface area were determined following 3-dimensional reconstruction using CTAn and CTVol software (Bruker Corporation).

### Metabolomics

At E14.5, maternal blood was collected by cardiac puncture and fetal blood was collected by pooling trunk blood from decapitated fetuses. Serum was separated using SST vacutainer tubes (Beckton Dickinson) and frozen at −80°C. Samples were prepared using the automated MicroLab STAR system (Hamilton Company) and analyzed on GC/MS, LC/MS and LC/MS/MS platforms by Metabolon, Inc. Organic aqueous solvents were used to perform serial extractions for protein fractions, concentrated using a TurboVap system (Zymark) and vacuum dried. For LC/MS and LC-MS/MS, samples were reconstituted in acidic or basic LC-compatible solvents containing > 11 injection standards and run on a Waters ACQUITY UPLC and Thermo-Finnigan LTQ mass spectrometer, with a linear ion-trap frontend and a Fourier transform ion cyclotron resonance mass spectrometer back-end. For GC/MS, samples were derivatized under dried nitrogen using bistrimethyl-silyl-trifluoroacetamide and analyzed on a Thermo-Finnigan Trace DSQ fast-scanning single-quadrupole mass spectrometer using electron impact ionization. Chemical entities were identified by comparison to metabolomic library entries of purified standards. Following log transformation and imputation with minimum observed values for each compound, data were analyzed using one-way ANOVA to test for group effects. P- and q-values were calculated based on two-way ANOVA contrasts. Principal components analysis was used to visualize variance distributions. Supervised Random Forest analysis was conducted to identify metabolomics prediction accuracies. All reference data is listed in Table S1.

#### Candidate metabolite selection for supplementation experiments

Physiologically relevant metabolite concentrations were determined based on reported murine serum concentrations from the mouse multiple tissue metabolomic database (MMMDB), human serum from the human metabolome database (HMDB), and existing literature. All reference literature is listed in Table S2.

### *In vivo* metabolite supplementation

To test the effects of the candidate microbiome-dependent fetal metabolites, the Metab cocktail or vehicle control was administered intraperitoneally once daily from E0.5 to E14.5 to minimize stress to pregnant dams. Metabolite dosages were calculated based on fetal serum metabolomic data and physiologically relevant metabolite concentrations reported in literature, to reflect the daily levels needed to match those observed in SPF fetal serum (Table S2). Metabolite concentrations were calculated based on physiological levels in mouse or human blood (Table S2), total blood volume of pregnant mouse dams (approximately 58.5 ml/kg (*7*)), and relative reductions observed in fetal sera of ABX fetuses compared to SPF fetuses (Table S2). The metabolite stock solution consisted of: 29.64 μM imidazole propionate, 714.096 μM N,N,N-trimethyl-5-aminovalerate, 13.452 μM 4-hydroxyphenylacetate, 316.008 μM phenol sulfate, 428.868 μM indolepropionate, 360.24 μM indoxyl glucuronide, 323.76 μM N-methylproline, 595.080 μM phenylacetylglycine, 957.6 μM trimethylamine N-oxide, 118.56 μM taurodeoxycholate, 160.8768 μM biotin, 893.76 μM hippurate, 42.864 μM 2-(4-hydroxyphenyl)propionate, 222.072 μM cinnamoylglycine, 273.6 nM equol glucuronide, 433.2 μM 2-aminophenol sulfate, 1.55952 mM 3-indoxyl sulfate and 268.128 μM p-cresol sulfate in 0.1 M PBS. The stock solution was then diluted 1:100 in sterile 0.1 M PBS, and 200 ul of the working solution was injected intraperitoneally into E0.5 ABX dams once a day daily for 14 days. For SCFA treatment, sodium propionate (25 mM), sodium butyrate (40 mM), and sodium acetate (67.5 mM) were supplemented to drinking water of pregnant ABX dams from E0-E14.5. These concentrations were determined based on existing studies demonstrating that these concentrations were able to sufficiently penetrate host tissues distal from the gut and restore circulating physiological concentrations (*8*). SCFA-supplemented drinking water and sodium-matched control drinking water were sterile-filtered and made fresh every 4 days. To assess placental phenotypes, dams treated with metabolites were sacrificed on E14.5, and placentas were harvested and processed as described in sections above.

### HUVEC angiogenesis assay

First-generation HUVECs (ThermoFisher Scientific, C0035C)) were passaged to third or fourth generations in Medium 200 (ThermoFisher Scientific) with Large Vessel Endothelial Supplement (LVES, ThermoFisher Scientific) and penicillin (10 U/ml)/streptomycin (10 μg/ml; Sigma-Aldrich) at 37°C, 5% CO_2_. Tube formation assays were performed using tissue-culture treated μslides (Ibidi). Briefly, 10 μl of ice-cold Geltrex LVES-free Reduced Growth Factor Basement Membrane Matrix (ThermoFisher Scientific) was added using pre-chilled pipettes to each well of a chilled μslide, and μslides were transferred to 37°C for 30 minutes for matrix polymerization following confirmation of equal matrix distribution across all wells. Third or fourth generation HUVECs were resuspended in LVES-free Medium 200 (minimal endothelial cell media) with penicillin (10 U/ml)/streptomycin (10 μg/ml; Sigma-Aldrich), and then resuspended with metabolite-supplemented media at the concentrations indicated in Table S3 at 2 x 10^5^ cells/ml, and 50 μl of cell suspension was added to each well. HUVECs in LVES-free Medium 200 were used as a negative control, and Medium 200 with 2%-FBS was used as a positive control. All conditions were performed in triplicate for each experiment, and each experiment was repeated 3 times. μslides were incubated at 37°C, 5% CO_2_ for 12 hours and imaged using a Leica DMi8 microscope. Image analysis and quantification was performed using Fiji (*9*) and the Angiogenesis analyzer plugin (*10*), and total branching distance values were averaged first for technical replicate by experiment, and then by biological replicate across multiple experiments, which is displayed in final figures.

### Statistical Methods

Statistical analysis was performed using Prism software (GraphPad). Data were assessed for normal distribution and plotted in the figures as mean ± SEM. For each figure, unless otherwise specified, “n” reflects the number of independent maternal biological replicates. Whole litters were only excluded if the total number of viable conceptuses was less than 4, and otherwise no conceptuses were excluded from any litters represented. Differences among 2 or more groups with only one variable were assessed using one-way ANOVA with Tukey *post hoc* test. Taxonomic comparisons from 16S rRNA gene sequencing analysis were analyzed by Kruskal-Wallis test with Tukey’s *post hoc* test. Differences between only 2 groups with one variable were assess using unpaired two-tailed t test. Two-way ANOVA with Tukey *post hoc* test was used for comparison of 2 or more groups with two variables. For all figures, significant differences emerging from the above tests are indicated by *p < 0.05, **p < 0.01, ***p < 0.001, ****p < 0.0001, and notable non-significant differences are indicated by “n.s.”.

### Data Availability

Data generated or analyzed during this study are included in this published article and its supplementary information files. The 16S rRNA gene sequencing data that support the findings of this study are available in Table S4 and have been deposited to the Qiita database with study ID 14433. Transcriptomic data that support the findings of this study are available in the Table S5 and have been deposited to the Gene Expression Omnibus (GEO) repository with accession number GSE198947.

## Supplemental Text

Results from this study reveal a role for the maternal microbiome in supporting placental growth and vascular development during pregnancy. We find that depletion of the maternal microbiome by oral treatment with antibiotics impairs placental development, which is prevented by supplementation with SCFAs derived from microbial fermentation of complex dietary carbohydrates, strongly suggesting that the maternal microbiome particularly in the gut is responsible. This distinction is notable in light of several studies reporting the existence of a microbiome within the placenta itself (*19–22*). Whether there are microbes native to the placental environment remains controversial, as additional evidence has suggested that microbial sequences detected in placental samples are due to laboratory contamination or reflect pathogenic entry into the human placenta, rather than a native placenta-associated microbiota (*23, 24*). In addition, evidence supporting the presence of viable symbiotic microbes in the placenta, as opposed to microbial genetic material, captured by immune cells for example, is lacking. In mice, the existence of live microbes in the placenta that reach the fetus is repudiated by the ability to generate germ-free animals by cesarean hysterectomy shortly before parturition. Findings in this study in mice provide proof-of-principle that the maternal gut microbiome and gut microbial metabolites are required for supporting placental morphogenesis.

We find that maternal supplementation with SCFAs increases placental size and feto-placental vasculature across two mouse models, which is not seen with maternal supplementation with a consortium of other key metabolites that are modulated by the maternal microbiome. SCFA concentrations increase with pregnancy (*25*), and reach fetal circulation via transplacental transport through monocarboxylate transporters (*26*). Consistent with our findings in a mouse model of maternal protein restriction, maternal consumption of high fat diet during pregnancy resulted in placental hypoxia and alterations in placental blood vessel structure, which were correlated with reductions in SCFA-producing bacteria in the maternal gut microbiota. Maternal supplementation with the SCFA butyrate prevented the placental impairments (*27*), supporting a role for SCFAs in promoting placental development. While the mechanisms underlying the influences of SCFAs on the placenta remain unknown, SCFA receptors are known to be expressed by uteroplacental tissues across mammals (*28*), suggesting the potential for direct receptor-mediated signaling of SCFAs to placental cells. Consistent with this, we show that select SCFAs increase angiogenic tube formation by HUVEC cells, which may explain the ability of maternal SCFA supplementation during pregnancy to increase feto-placental vasculature *in vivo*.

Beyond interactions with the placenta, SCFAs have been reported to support the development of fetal intestinal, pancreatic, and neural tissues in mice (*12*). Reduced concentrations of the SCFA acetate have been reported in women with preeclampsia, and perinatal acetate supplementation is sufficient to promote fetal thymic CD4^+^ T cell and Treg development in germ-free mice (*29*). Whether effects of SCFAs on placental structure and function may contribute to the reported influences of SCFAs on fetal development remains unclear.

Placental insufficiencies, including vascular impairments, have long been linked to adverse outcomes in developing offspring and increased risk for chronic disease during adulthood (*30*). Placental vascular deficits, as indicated by elevated umbilical artery resistance *in utero*, are associated with reduced fetal weight and length, as well as reduced body mass, elevated systolic blood pressure, and reduced left ventricular mass during childhood (*31*). Consistent with this, reduced placental size is correlated with increased incidence of hypertension during adulthood (*30*). Moreover, placental vascular deficiencies characteristic of hypertensive disorders of pregnancy, and preeclampsia in particular, are associated with increased risk for myriad diseases in adulthood, including cardiovascular disease, kidney disease, and cognitive impairment (*32*). Findings from our study reveal that metabolic functions contributed by the maternal gut microbiome during pregnancy are integral to supporting placental growth and vascularization in mice. Advancing our understanding of how the maternal gut microbiome impacts placental structure and function may inform new approaches to promote maternal and fetal health and to decrease risk for chronic diseases.

**Fig. S1:**
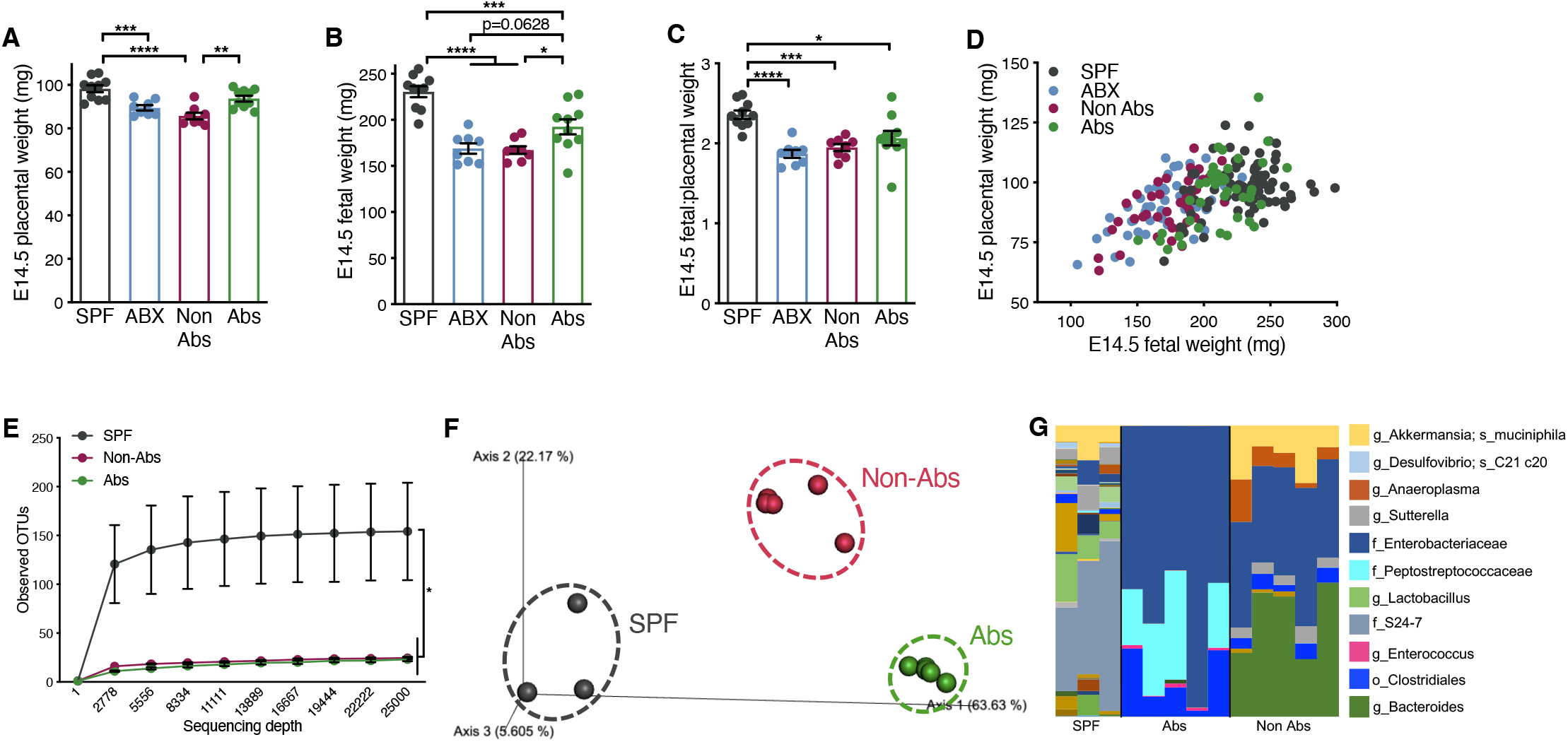
Antibiotic-induced reductions in placental and fetal weight are not due to off-target effects of absorbable antibiotics. **(A)** E14.5 placental weights by litter average (SPF (*n* = 10; same as shown in Fig. 1), ABX (ampicillin, metronidazole, neomycin and vancomycin; *n* = 9), Nonabsorbable antibiotics (Non-Abs; neomycin and vancomycin; *n* = 8) and absorbable antibiotics (Abs; ampicillin and metronidazole; *n* = 10). (**B)** E14.5 fetal weights by litter average from litters shown in Fig. S1A. (**C)** E14.5 fetal to placental weight ratios for litters shown in Fig. S1A-B. (**D)** Linear correlation of E14.5 placental to fetal weight ratios for all individual conceptuses (SPF (*n* = 80), ABX (*n* = 49), Non-Abs (*n* = 40) and Abs (*n* = 42)) from litters shown in Fig. S1A-C. **(E)** Alpha-rarefaction curves of 16S rRNA gene sequencing results from E14.5 maternal fecal samples collected from independent cages (SPF (*n* = 3), Non-Abs (*n* = 5) and Abs (*n* = 5)). (**F)** Weighted UniFrac analysis of 16S rRNA gene sequencing results from E14.5 maternal fecal samples shown in Fig. S1E. (**G)** Taxa bar plots for 16S rRNA gene sequencing results from E14.5 maternal fecal samples shown in Fig. S1E-F. Data represent mean ± SEM; statistics were performed with one-way ANOVA with Tukey *post hoc* test. \**P* < 0.05; \*\**P* < 0.01; \*\*\**P* < 0.001; \*\*\*\**P* < 0.0001.

**Fig. S2:**
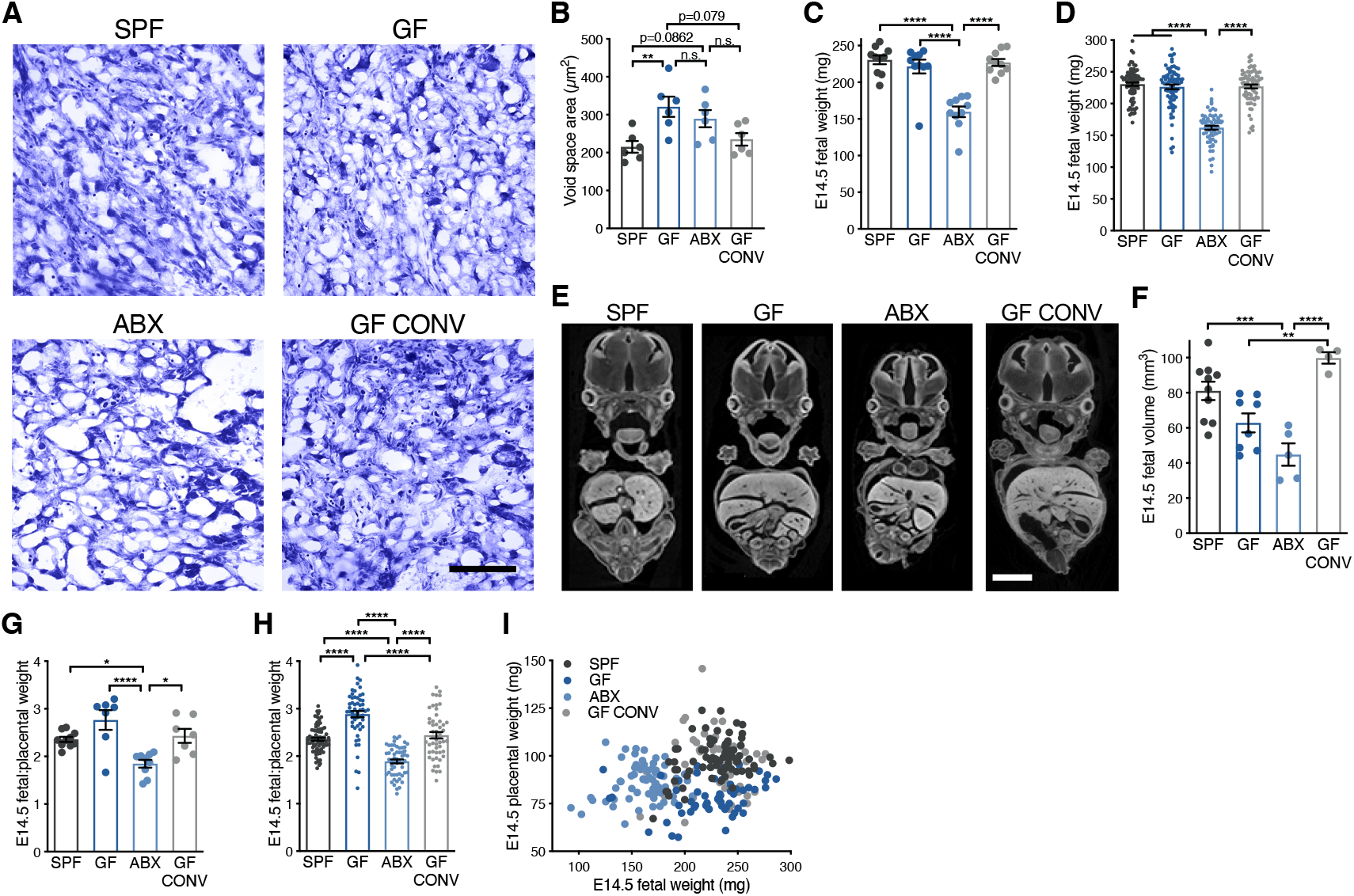
Maternal microbiome depletion differentially affects E14.5 fetal weight. **(A)** Representative images of thionin-stained E14.5 placental labyrinth tissues from SPF, GF, ABX and GF CONV litters (scale bar = 250 μm)**. (B)** Quantification of average placental labyrinth void space area from litters shown in Fig. S2A (SPF (*n* = 6), GF (*n* = 6), ABX (*n* = 6) and GF CONV (*n* = 6)). (**C)** E14.5 fetal weights by litter average (SPF (*n* = 10), GF (*n* = 10), ABX (*n* = 10) and GF CONV (*n* = 10). (**D)** E14.5 fetal weights for each individual from litters shown in Fig. S2C (SPF (*n* = 80), GF (*n* = 81), ABX (*n* = 65) and GF CONV (*n* = 77) litters. (**E)** Representative cross-sections of E14.5 whole fetal *μ*CT reconstructions from SPF, GF, ABX and GF CONV litters (scale = 1mm). (**F)** Quantification of E14.5 whole fetal volumes from *μ*CT reconstructions by litter average (SPF (*n* = 10), GF (*n* = 8), ABX (*n* = 5) and GF CONV (*n* = 4). (**G)** E14.5 fetal to placental weight ratio for litters shown in Fig. S2C (SPF (*n* = 10), GF (*n* = 10), ABX (*n* = 10) and GF CONV (*n* = 10). **(H)** E14.5 fetal to placental weight ratios for individual conceptuses from litters shown in Fig. S2G (SPF (*n* = 80), GF (*n* = 81), ABX (*n* = 65) and GF CONV (*n* = 77) litters. **(I)** Linear correlation of E14.5 placental to fetal weight ratios for individual conceptuses shown in Fig. S2H. Data represent mean ± SEM; statistics were performed with one-way ANOVA with Tukey *post hoc* test. \**P* < 0.05; \*\**P* < 0.01; \*\*\**P* < 0.001; \*\*\*\**P* < 0.0001.

**Fig. S3:**
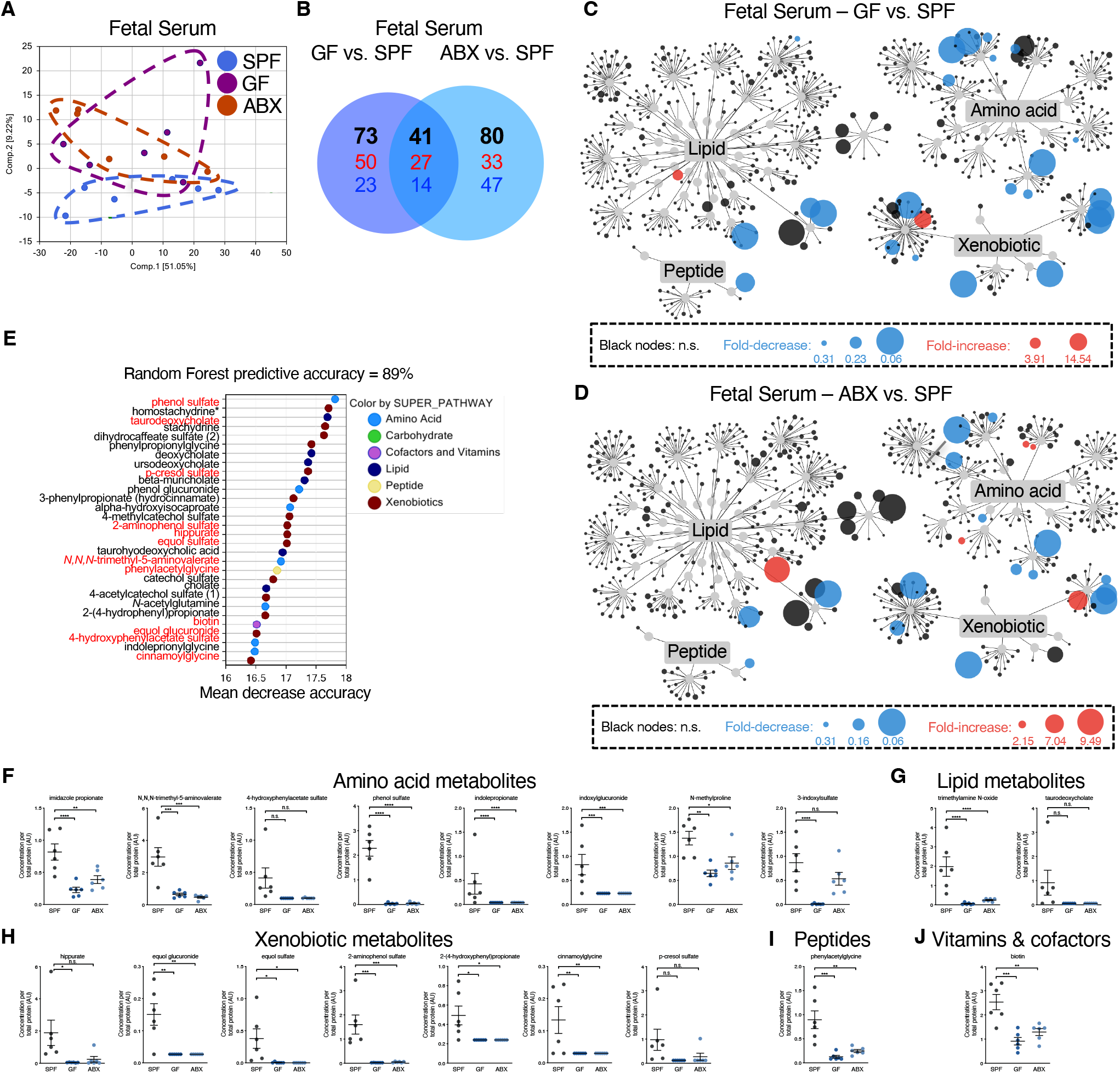
The maternal microbiome modulates the fetal serum metabolome. **(A)** Principal component analyses of E14.5 fetal serum (SPF (*n* = 6), GF (*n* = 6) and ABX (*n* = 6)). (**B)** Significantly altered metabolites (black text), both downregulated (red text) and upregulated (blue text) in litters represented in Fig. S3B. (**C)** Global metabolite profile changes in GF fetal serum relative to SPF fetal serum. (**D)** Global metabolite profile changes in ABX fetal serum relative to SPF fetal serum. **(E)** Random Forest analysis (predictive accuracy of 89%) identifying the top 30 fetal serum metabolites that distinguish samples as GF or ABX, relative to SPF (metabolites used for *in vivo* supplementation labeled with red text). (**F)** Relative concentrations of amino acid metabolites used for *in vivo* metabolite supplementation from litters represented in Fig. S3A-B. (**G)** Relative concentrations of lipid metabolites used for *in vivo* metabolite supplementation from litters represented in Fig. S3A-B. (**H)** Relative concentrations of xenobiotic metabolites used for *in vivo* metabolite supplementation from litters represented in Fig. S3A-B. (**I)** Relative concentration of peptide metabolites used for *in vivo* metabolite supplementation from litters represented in Fig. S3A-B. (**J)** Relative concentration of vitamin & cofactor metabolites used for *in vivo* metabolite supplementation from litters represented in Fig. S3A-B. Data represent mean ± SEM; statistics were performed with one-way ANOVA with Tukey *post hoc* test. \**P* < 0.05; \*\**P* < 0.01; \*\*\**P* < 0.001; \*\*\*\**P* < 0.0001.

**Fig. S4:**
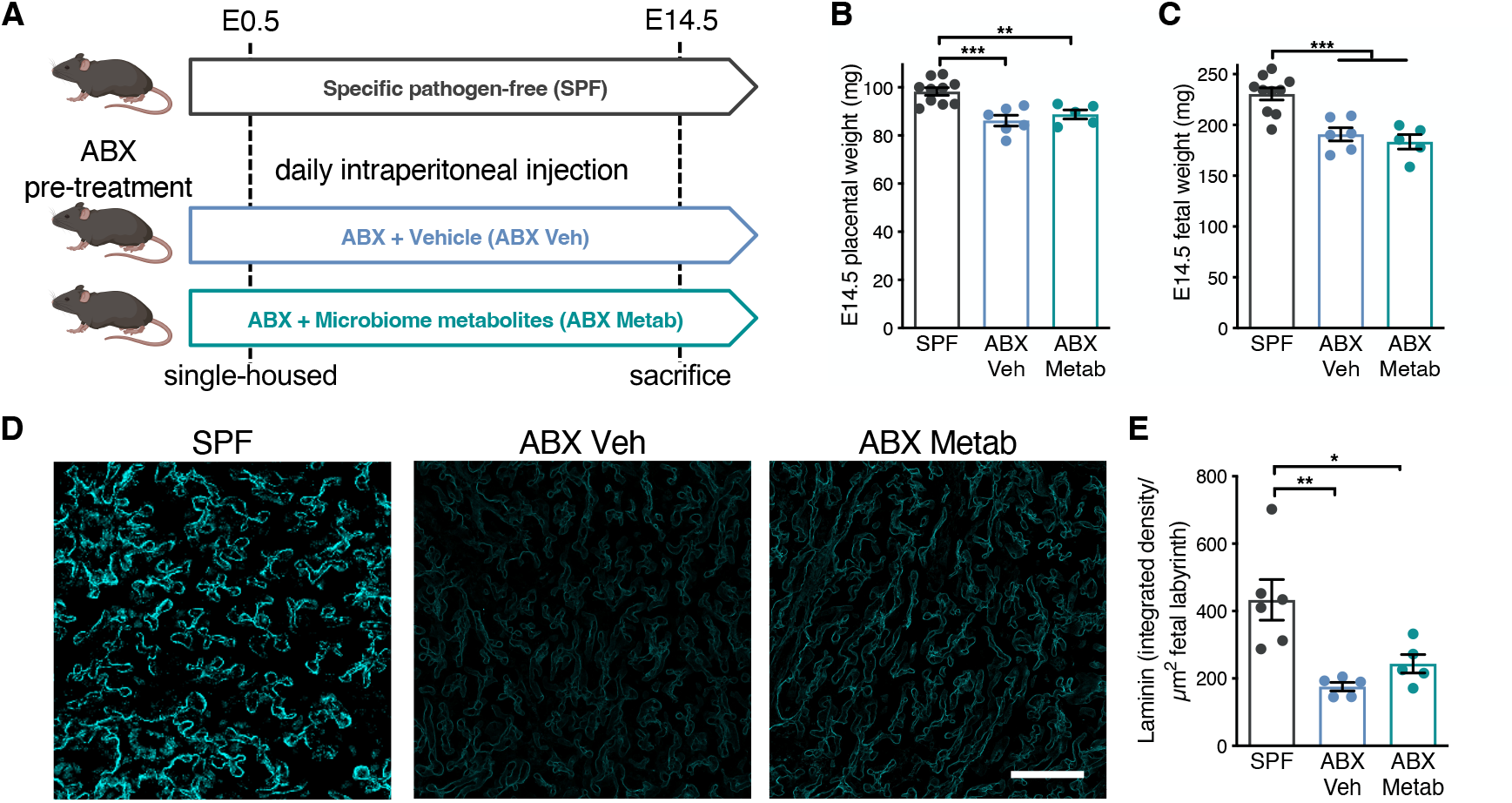
Select microbiome-dependent metabolites do not rescue placental growth and vascular impairments induced by maternal microbiome depletion. **(A)** Schematic for microbiome-dependent metabolite supplementation in ABX mice. (**B)** E14.5 placental weights by litter average (SPF (*n* = 10; same as shown in Fig. 1A), ABX Veh (*n* = 6) and ABX Metab (*n* = 5)). (**C)** E14.5 fetal weights for litter averages shown in Fig. S4B. (**D)** Representative images of laminin-stained E14.5 placental labyrinth from SPF, ABX Veh and ABX Metab litters. (**E)** Quantification of raw integrated density of laminin intensity, normalized to fetal labyrinth total area (SPF (*n* = 6; same as shown in Fig. 2C), ABX Veh (*n* = 5) and ABX Metab (*n* = 5)). Data represent mean ± SEM; statistics were performed with one-way ANOVA with Tukey *post hoc* test. \**P* < 0.05; \*\**P* < 0.01; \*\*\**P* < 0.001; \*\*\*\**P* < 0.0001.

**Fig. S5:**
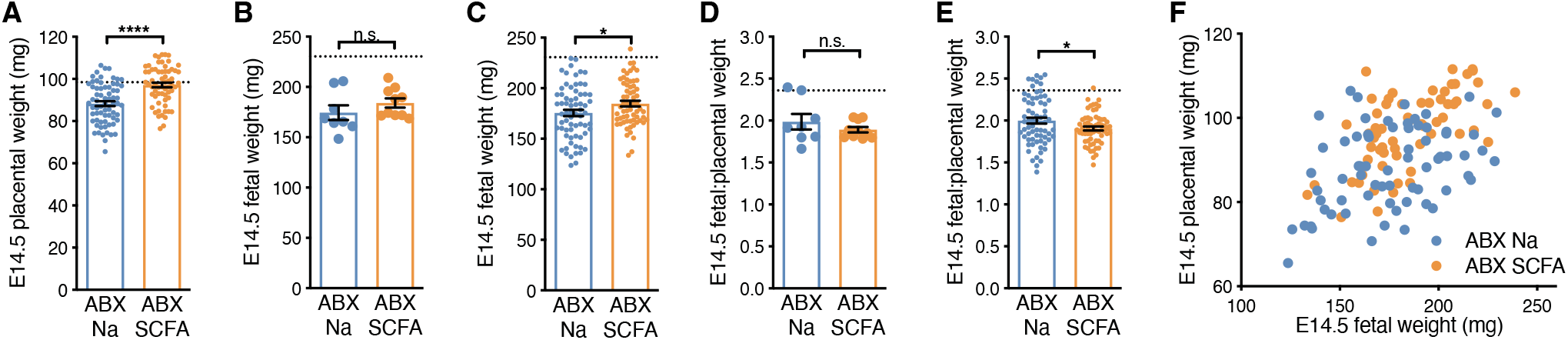
SCFA supplementation does not alter E14.5 fetal growth in microbiota-depleted dams. **A)** E14.5 placental weights for each individual from litters shown in Fig. 3A (ABX Na (*n* = 68) and ABX SCFA (*n* = 63); hashed line represents SPF litter average value shown in Fig. 1B). **B)** E14.5 fetal weights by litter average for litters shown in Fig. S5A (hashed line represents SPF litter average value shown in Fig. S2B). **C)** E14.5 fetal weights for each individual from litters shown in Fig. S5A (hashed line represents SPF litter average value shown in Fig. S2C). **D)** E14.5 fetal to placental weight ratios by litter average for litters shown in Fig. S5A-C (hashed line represents SPF litter average value shown in Fig. S2F). **E)** E14.5 individual fetal to placental weight ratios from litters shown in Fig. S5D (hashed line represents SPF litter average value shown in Fig. S2G). **F)** Linear correlation of E14.5 placental to fetal weight ratios for individual conceptuses shown in Fig. S5A-C. Means ± SEM (error bars) plotted; statistics were performed with student’s t test. *\*P* < 0.05; \*\**P* < 0.01; \*\*\**P* < 0.001; \*\*\*\**P* < 0.0001.

**Fig. S6:**
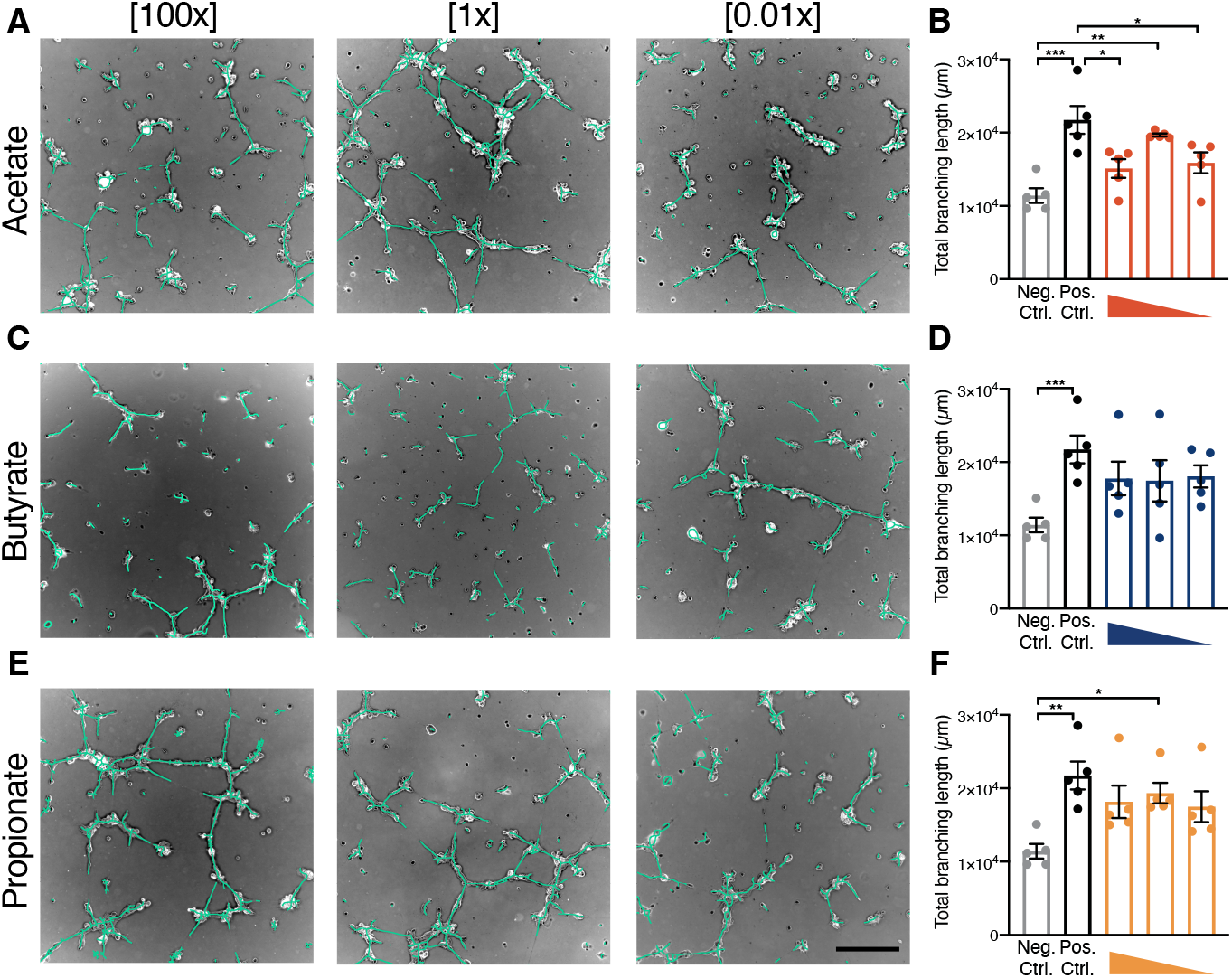
SCFAs promote HUVEC tube formation *in vitro*. **(A)** Representative images for HUVECs treated with 4 mM, 40 *μM* and 0.4 μM acetate, respectively. (**B)** Quantification of acetate-supplemented HUVEC tube formation assays. (**C)** Representative images for HUVECs treated with 500 μM, 5 μM and 0.05 μM butyrate, respectively. (**D)** Quantification of propionate-supplemented HUVEC tube formation assays. (**E)** Representative images for HUVECs treated with 500 μM, 5 μM and 0.05 μM propionate, respectively. (**F)** Quantification of butyrate-supplemented HUVEC tube formation assays. Scale bar = 250 μm. Data represent mean ± SEM; statistics were performed with one-way ANOVA with Tukey *post hoc* test. \**P* < 0.05; \*\**P* < 0.01; \*\*\**P* < 0.001; \*\*\*\**P* < 0.0001.

**Fig. S7:**
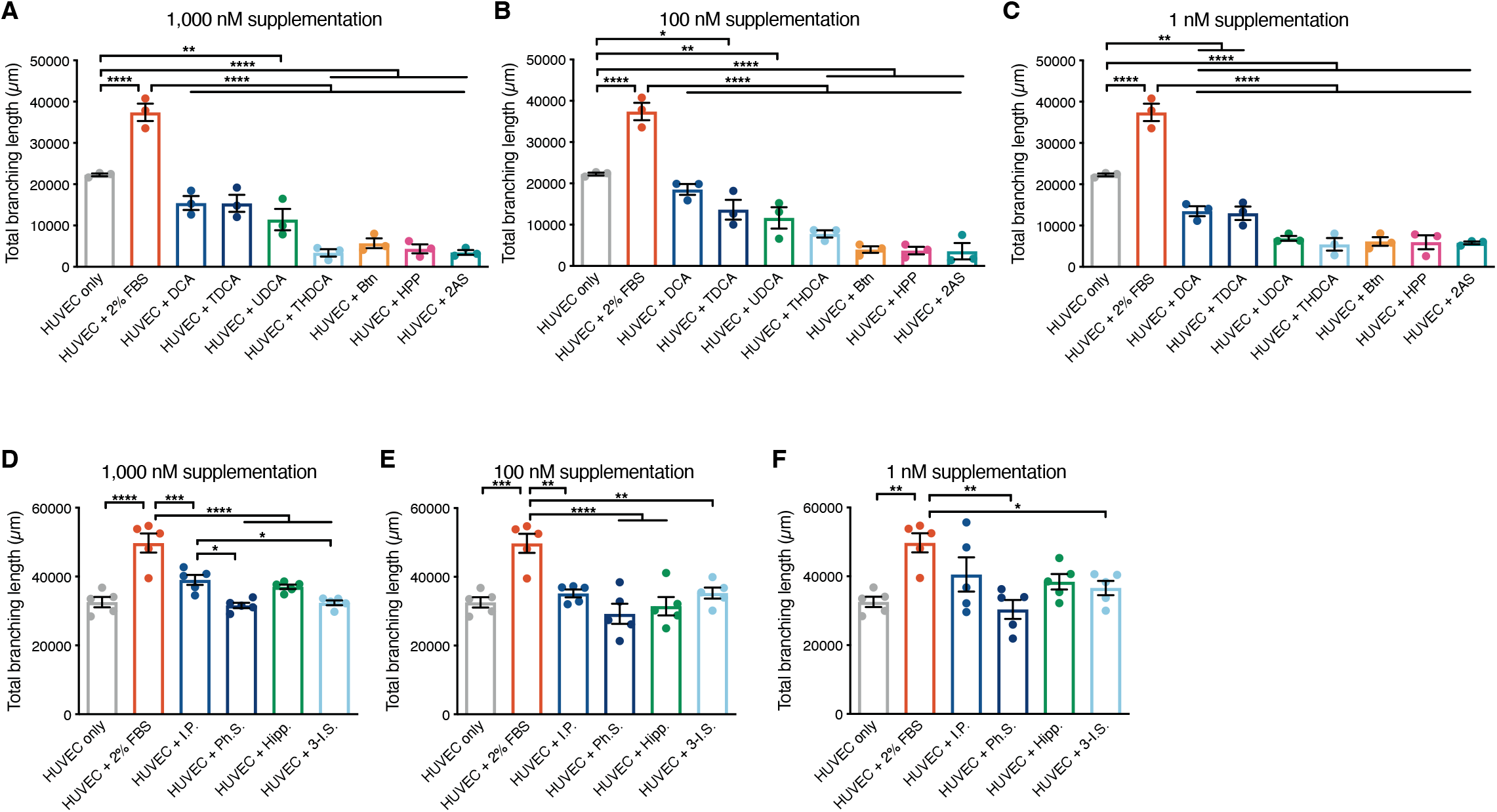
Select microbiome-dependent metabolites do not promote HUVEC tube formation *in vitro*. **(A)** Quantification of HUVEC branching length in non-supplemented media (HUVEC only, negative control), 2% FBS-supplemented media (positive control), and supplementation with 1,000 nM deoxycholate (DCA), taurodeoxycholate (TDCA), ursodeoxycholate (UDCA), taurohyodeoxycholate (THDCA), biotin (Btn), 2-(4-hydrophenyl)propionate (HPP) and 2-aminophenol sulfate (2AS), respectively. (**B)** Quantification of HUVEC branching length in nonsupplemented media, 2% FBS-supplemented media, and supplementation with 100 nM DCA, TDCA, UDCA, THDCA, Btn, HPP and 2AS, respectively. (**C)** Quantification of HUVEC branching length in non-supplemented media, 2% FBS-supplemented media, and supplementation with 1 nM DCA, TDCA, UDCA, THDCA, Btn, HPP and 2AS, respectively. (**D)** Quantification of HUVEC branching length in non-supplemented media, 2% FBS-supplemented media, and supplementation with 1,000 nM imidazole propionate (I.P.), phenol sulfate (Ph.S.), hippurate (Hipp.) and 3-indoxyl sulfate (3-I.S.), respectively. (**E)** Quantification of HUVEC branching length in non-supplemented media, 2% FBS-supplemented media, and supplementation with 100 nM imidazole propionate (I.P.), phenol sulfate (Ph.S.), hippurate (Hipp.) and 3-indoxyl sulfate (3-I.S.), respectively. (**F)** Quantification of HUVEC branching length in non-supplemented media, 2% FBS-supplemented media, and supplementation with 1 nM imidazole propionate (I.P.), phenol sulfate (Ph.S.), hippurate (Hipp.) and 3-indoxyl sulfate (3-I.S.), respectively. Data represent mean ± SEM; statistics were performed with one-way ANOVA with Tukey *post hoc* test. \**P* < 0.05; \*\**P* < 0.01; \*\*\**P* < 0.001; \*\*\*\**P* < 0.0001.

**Fig. S8:**
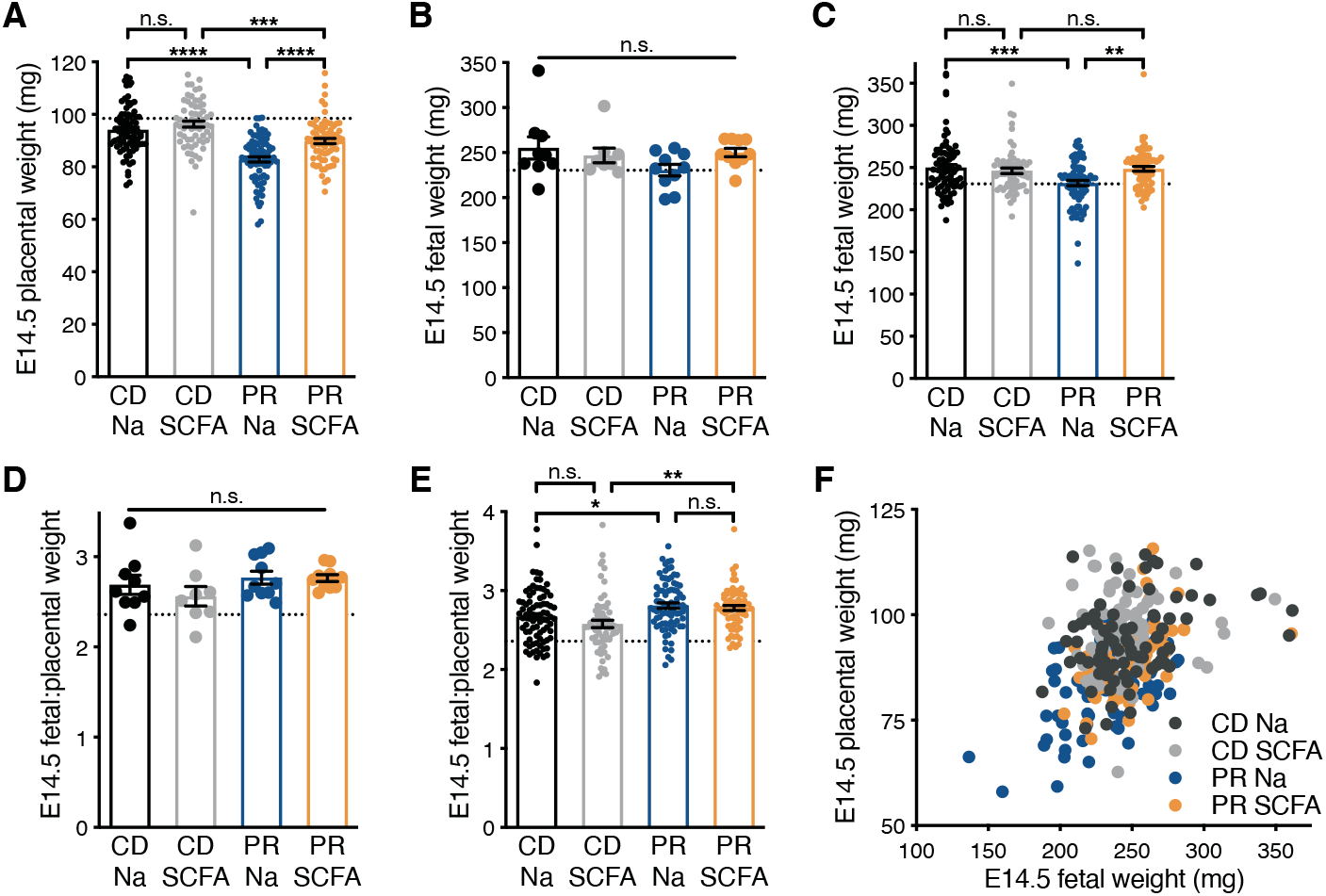
Maternal SCFA supplementation modestly increases E14.5 fetal growth in protein-restricted dams. **(A)** E14.5 placental weights for each individual from litters shown in Fig. 4A (CD Na (*n* = 80), CD SCFA (*n* = 65), PR Na (*n* = 81) and PR SCFA (*n* = 72); hashed line represents SPF litter average value shown in Fig. 1B). (**B)** E14.5 fetal weights by litter average for litters shown in Fig. S8A (hashed line represents SPF litter average value shown in Fig. S2B). (**C)** E14.5 fetal weights for each individual from litters shown in Fig. S8A (hashed line represents SPF litter average value shown in Fig. S2C). (**D)** E14.5 fetal to placental weight ratios by litter average for litters shown in Fig. S5A-C (hashed line represents SPF litter average value shown in Fig. S2F). (**E)** E14.5 individual fetal to placental weight ratios from litters shown in Fig. S5D (hashed line represents SPF litter average value shown in Fig. S2G). (**F)** Linear correlation of E14.5 placental to fetal weight ratios for individual conceptuses shown in Fig. S5A-C. Data represent mean ± SEM; statistics were performed with two-way ANOVA with Tukey *post hoc* test. \**P* < 0.05; \*\**P* < 0.01; \*\*\**P* < 0.001; \*\*\*\**P* < 0.0001.

